# Perceptual Expectations do not Modulate Image Repetition Effects as Measured by Event-Related Potentials

**DOI:** 10.1101/132621

**Authors:** Daniel Feuerriegel, Owen Churches, Scott Coussens, Hannah A. D. Keage

**Affiliations:** School of Psychology, Social Work and Social Policy, University of South Australia; Melbourne School of Psychological Sciences, The University of Melbourne; School of Psychology, Flinders University

**Author notes:** Corresponding Author: Daniel Feuerriegel, Postal Address: Daniel Feuerriegel, Melbourne School of Psychological Sciences, Redmond Barry Building, The University of Melbourne, Parkville, Victoria, Australia.

**Keywords:** Repetition Suppression, Stimulus Specific Adaptation, Expectation, Prediction, ERP

## Abstract

Repeated stimulus presentation leads to complex changes in cortical neuron response properties, commonly known as repetition suppression or stimulus-specific adaptation. Circuit-based models of repetition suppression provide a framework for investigating patterns of repetition effects that propagate through cortical hierarchies. To further develop such models it is critical to determine whether (and if so, when) repetition effects are modulated by top-down influences, such as those related to perceptual expectation. We investigated this by presenting pairs of repeated and alternating face images, and orthogonally manipulating expectations regarding the likelihood of stimulus repetition. Event-related potentials (ERPs) were recorded from n=39 healthy adults, to map the spatiotemporal progression of stimulus repetition and expectation effects, and interactions between these factors, using mass univariate analyses. We also tested whether the ability to predict unrepeated (compared to repeated) face identities could influence the magnitude of observed repetition effects, by presenting separate blocks with predictable and unpredictable alternating faces. Multiple repetition and expectation effects were identified between 99-800ms from stimulus onset, which did not statistically interact at any point. Repetition effects in blocks with predictable alternating faces were smaller than in unpredictable alternating face blocks between 117-179ms and 506-652ms, and larger between 246-428ms. ERP repetition effects appear not to be modulated by perceptual expectations, supporting separate mechanisms for repetition and expectation suppression. However, previous studies that aimed to test for repetition effects, in which the repeated (but not unrepeated) stimulus was predictable, are likely to have conflated repetition and stimulus predictability effects.

**Highlights:**

- ERP face image repetition effects were apparent between 99-800ms from stimulus onset
- Expectations of stimulus image properties did not modulate face repetition effects
- The predictability of unrepeated stimuli influenced repetition effect magnitudes

## 1. Introduction

Cortical neurons adapt to dynamic and static features of an organism’s environment. Repeated exposure to a stimulus leads to changes in response properties of cortical neurons, known as repetition suppression or stimulus-specific adaptation (Desimone, 1996; Movshon & Lennie, 1979). Repetition suppression refers to a stimulus-specific reduction in a recorded signal of neuronal activity (e.g. firing rate, local field potential amplitude or fMRI BOLD signal change) to repeated compared to unrepeated stimuli (for reviews see Grill-Spector, Henson & Martin, 2006; Kohn, 2007; Vogels, 2016) and is one phenomenon in a wider family of adaptation effects, which includes response increases with stimulus repetition (reviewed in Segaert et al., 2013) and exposure-dependent changes in stimulus selectivity (e.g. Sawamura et al., 2006; Wissig & Kohn, 2012).

Theories of perception based on hierarchically-organised predictive coding (Rao & Ballard, 1999; Friston, 2005) conceptualise repetition suppression as a reduction of prediction error signals due to fulfilled perceptual expectations weighted toward recently-encountered stimuli (e.g. Summerfield et al., 2008; Auksztulewicz & Friston, 2016). Repetition suppression is proposed to occur via lateral or feedback inhibition from neurons that actively generate predictions within hierarchically-organised sensory systems (Friston, 2005). It is unclear whether predictive coding models can account for the vast range of adaptation effects reported in the literature (reviewed in Solomon & Kohn, 2014; Whitmire & Stanley, 2016; Kaliukhovich & Vogels, 2016). However these models, and circuit-based models in general, provide a framework for investigating repetition effects that propagate through cortical hierarchies via feedforward, feedback and lateral projections (e.g. Dhruv & Carrandini, 2014; Larsson et al., 2016; Malmiciera et al., 2015; Wissig & Kohn, 2012; Patterson et al., 2013).

To test and develop predictive coding models of repetition effects it is critical to identify how stimulus repetition effects relate to other forms of perceptual expectation, which may be separable or interactive within sensory systems (Grotheer & Kovacs, 2016). An influential study of Summerfield et al. (2008) investigated relationships between stimulus repetition and perceptual expectations based on the likelihood that a certain stimulus (or sequence of stimuli) would appear. They presented pairs of repeated and alternating faces in blocks with high and low proportions of repetition trials, and reported that BOLD repetition effects in the fusiform face area (FFA; Kanwisher et al., 1997) were larger in high repetition probability blocks. These results were interpreted as a modulation of repetition suppression by perceptual expectation. The findings of Summerfield et al. have been replicated several times (Kovacs et al., 2012, 2013; de Gardelle et al., 2013; Grotheer & Kovacs, 2014; Choi et al., 2017) although these expectation effects appear to be attention-dependent (Larsson & Smith, 2012) and restricted to highly familiar stimulus categories (Kovacs et al., 2013; Grotheer & Kovacs, 2014).

The repetition by block interactions in Summerfield et al. (2008) and subsequent replications may have actually identified additive effects of stimulus repetition and expectation. Responses to expected (high occurrence probability) repetitions were compared with surprising (low occurrence probability) alternations in high repetition probability blocks, and surprising repetitions with expected alternations in low probability blocks. Expected stimuli evoke smaller BOLD signals in FFA than surprising stimuli (den Ouden et al., 2010; Egner et al., 2010), and so larger repetition effects in high repetition probability blocks are likely a result of additive effects of stimulus repetition and expectation. More recent studies that independently manipulated repetition and expectation have reported mixed results. Grotheer and Kovacs (2015) found additive effects that were separable in time. However, Amado et al. (2016) reported that repetition effects were larger for surprising stimuli, driven by large surprise-related BOLD signal increases only for alternating stimuli. This pattern is also visible in results of many earlier studies (Kovacs et al., 2012; de Gardelle et al., 2013; Larsson & Smith, 2012; Grotheer et al., 2014; Choi et al., 2017; reviewed in Kovacs & Vogels, 2014).

Surprise effects that preferentially affect responses to alternating stimuli indicate that stimulus repetition may actually suppress the expression of prediction errors arising from violated perceptual expectations. However, these results might also be due to the experimental trial structure in the studies mentioned above. In these experiments adapter and test stimuli (i.e. the first and second stimuli presented in each trial) were the same stimulus image in repetition trials. In these trials the image properties of the repeated stimulus could be predicted after viewing the adapter stimulus. However, the alternating test stimulus was randomly-chosen from the stimulus set, and so image-specific expectations could not be formed for alternating test stimuli. Surprise-related signal increases for alternating stimuli may have arisen from violations of image-specific expectations of repeated stimuli. However, no image-specific expectations regarding alternating stimuli could be formed or violated, preventing large signal increases for surprising repetitions.

Additionally, it is possible that repetition effects observed in these widely-used stimulus repetition designs are caused or enhanced by imbalances of stimulus expectations available for repeated and alternating stimuli (Feuerriegel, 2016; Pajani et al., 2017). In this study we define the ability to form image-specific expectations about an upcoming stimulus as the predictability of that stimulus.

It is also unclear whether larger repetition effects for surprising stimuli are specific to the BOLD signal, or whether they can also be identified in electrophysiological recordings. Microelectrode studies of macaque inferior temporal neurons did not report interactions between stimulus repetition and expectation for firing rates or local field potentials (Kaliukhovich & Vogels, 2011; 2014). The only ERP investigation of these effects that presented visual stimuli (Summerfield et al., 2011) used an analysis design which confounded stimulus repetition and expectation, as described above. Electrophysiological evidence is especially important for evaluating predictive coding models, as prediction error signals are hypothesised to be generated by superficial pyramidal cells that contribute to scalp-recorded EEG responses (Friston & Kiebel, 2009; Auksztulewicz & Friston, 2016).

We investigated relationships between stimulus repetition effects and perceptual expectations by presenting repeated and alternating face pairs in high and low repetition probability contexts. In addition, separate experimental blocks were run in which alternating face stimuli were either predictable or unpredictable. Using mass univariate analyses of ERPs we were able to map the spatiotemporal patterns of repetition and expectation effects. We aimed to identify latencies from stimulus onset at which repetition effects are modulated by perceptual expectations, or by the ability to predict the alternating stimulus image. By allowing image-specific expectations for alternating faces we were also able to test whether the large surprise-related responses for alternating stimuli (e.g. Amado et al., 2016) could be found for repeated stimuli (i.e. when image-specific expectations for alternating stimuli are violated). We expected to find larger ERP repetition effects for surprising stimuli, consistent with existing fMRI evidence (e.g. Amado et al., 2016; de Gardelle et al., 2013a), and larger ERP repetition effects in experimental blocks with unpredictable alternating faces.

## 2. Methods

### 2.1 Participants

Thirty-nine people participated in this experiment (10 males, mean age 25.3 ± 5.5, age range 18-38). All participants were native English speakers and had normal or corrected-to-normal vision, and were right-handed as assessed by the Flinders Handedness Survey (Nicholls et al., 2013). Three participants were excluded from analyses due to excessively noisy data. This experiment was approved by the Human Research Ethics committee of the University of South Australia.

### 2.2 Stimuli

Examples of stimuli are shown in Figure 1A. Ninety frontal images of faces (45 male, 45 female, neutral expression) were taken from the NimStim face database (Tottenham et al., 2009) and the Minear and Park Ebner face set (Minear & Park, 2004; Ebner, 2008). Images were converted to greyscale and cropped, resized and aligned so that the nose was in the horizontal center of the image and eyes of each face were vertically aligned. Image backgrounds were equated across Minear and Park and NimStim faces. Images were resized so that at a viewing distance of 60cm stimuli subtended approximately 3.15**°** x 4.30**°** of visual angle (135 x 180 pixels). Test stimulus images were created 20% larger than adapters. The SHINE toolbox (Willenbockel et al., 2010) was used to equate mean pixel intensity and contrast across images (Mean normalised pixel intensity = 0.38, mean RMS contrast = 0.12). Stimuli were presented against a grey background (normalised pixel intensity = 0.44).

**Figure 1.**
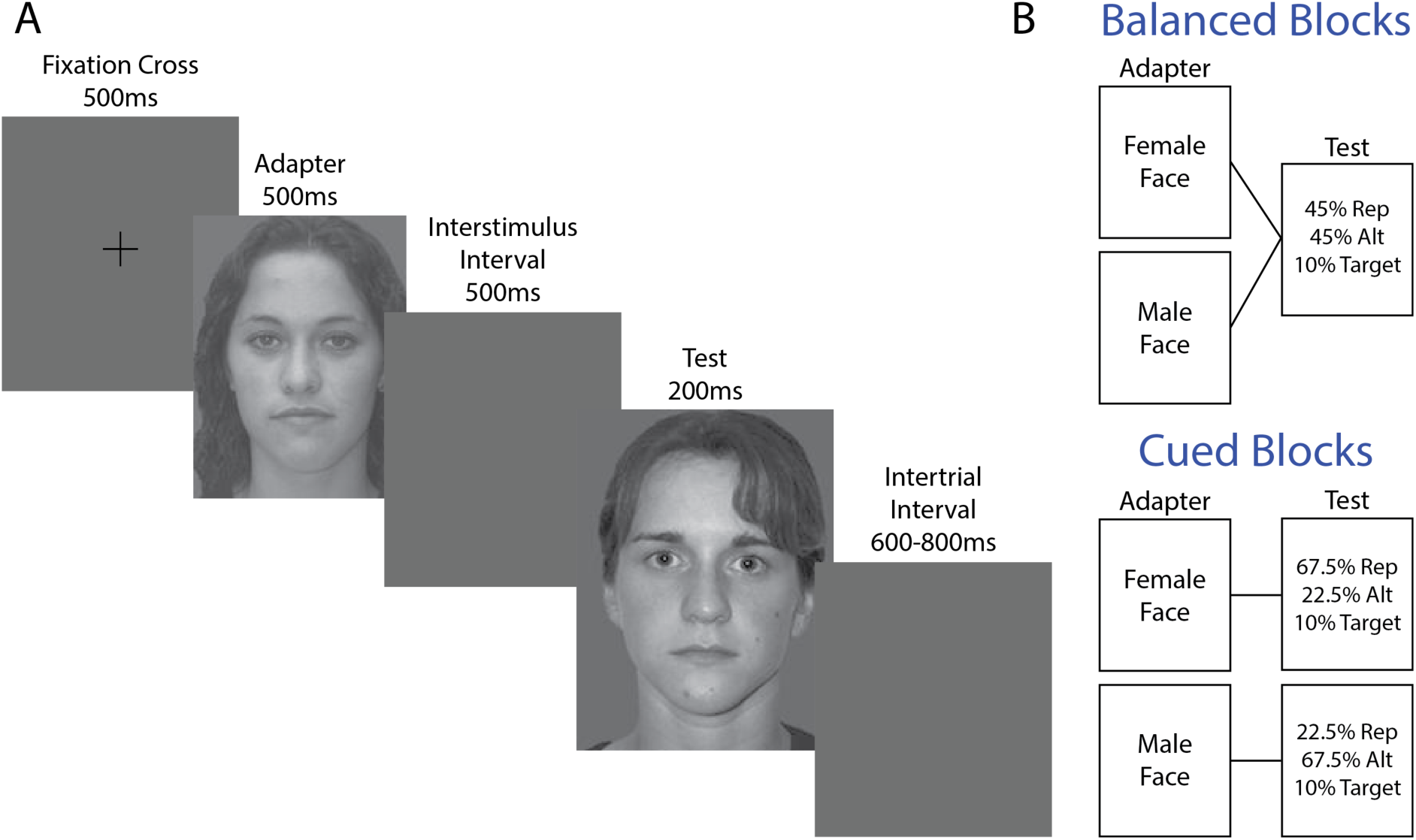
Trial diagram and repetition probability cueing design. A) In each trial adapter and test stimuli were presented separated by a 500ms interstimulus interval. Test stimuli were 20% larger than adapters. Shown is an example of an alternating trial in which adapters and tests are different face identities. B) Examples of experimental block types. In balanced blocks the probability of stimulus repetition is 45%. In cued blocks the probability of stimulus repetition varied by the gender of the adapter face. In this example a female adapter cues a high (67.5%) probability of stimulus repetition, whereas a male adapter cues a low (22.5%) probability of stimulus repetition. (two-column fitting image)

### 2.3 Procedure

Participants sat in a well-lit testing room 60cm in front of an LED monitor (refresh rate 60Hz). Stimuli were presented using custom scripts written in MATLAB (Mathworks) using functions from PsychToolbox v3.0.11 (Brainard, 1997; Kleiner et al., 2008). Behavioural responses were recorded using a one-button response box connected to the EEG amplifier.

In each trial face stimuli were presented as adapters (500ms) and tests (200ms) separated by a 500ms interstimulus interval (ISI) as shown in Figure 1A. Adapter faces were preceded by a fixation cross for 500ms. The intertrial interval (including fixation cross presentation duration) varied pseudorandomly between 1100-1300ms. Adapter and test stimuli were either the same face image (repetition trial) or two different faces of the same gender (alternation trials).

In target trials the test face was inverted along the horizontal plane (as done by Summerfield et al., 2008), and could be a repeated or alternating face with equal probability. Upon seeing a target participants were instructed to press a button on the response box as quickly as possible (response hands counterbalanced across participants). Responses between 200-1000ms from test stimulus onset were counted as correct responses. Target trials were presented as 10% of all trials.

There were 10 experimental blocks: 2 balanced blocks and 8 cued blocks (Figure 1B). In balanced blocks 45% of trials were repetition trials (neutral repetitions) and 45% were alternation trials (neutral alternations). The remaining 10% were target trials (5% repeated face targets, 5% alternating face targets). In cued blocks the probability of repetition was determined by the adapter face gender (overall probability of repetition 45%). As an example, for one participant female adapter faces signalled a high (67.5%) probability of stimulus repetition (expected repetitions) and a low (22.5%) probability of an alternation (surprising alternations). For the same participant male adapters signalled a low probability of repetition (surprising repetitions) and a high probability of an alternating trial (expected alternations). Gender cueing assignments were counterbalanced across participants. Balanced blocks were always presented before cued blocks to avoid carry-over effects of expectations from cued blocks.

Balanced and cued blocks were further subdivided into two block types, which differed with respect to the alternating test face identities. In both block types 4 faces (2 male, 2 female) were presented as adapters and repeated test stimuli (different faces used in each block). In repetition trials one of these faces appeared as both the adapter and test, with the restriction that the adapter in one trial could never be the test face in the preceding trial. In the first block type (AB blocks) alternating test faces were the other face identity of the same gender as the adapter. For example, if female face A was presented as the adapter then the test face could either be a repeat (AA) or the other female face (AB). In the second block type (AX blocks) the alternating face image was chosen randomly from a separate set of 46 faces (23 male, 23 female), with the restriction that each of the 23 faces within a gender set must be presented once before any face identity could be shown again. Adapters and tests were of the same gender in each trial. For example, if female face A was presented as the adapter, the test face could either be a repeat (AA) or a different randomly-selected female face (AX). AB and AX blocks alternated throughout the experiment (block order counterbalanced across participants). Face images used for adapter and repeated test stimuli allocated to each block were also counterbalanced across participants. A practice block was also presented before the experiment (AB block, 24 trials, using a separate set of 4 faces).

A total of 1600 nontarget trials and 180 target trials were presented. In balanced blocks there were 80 nontarget trials for each repeated and alternating condition. In cued blocks there were 240 nontarget trials for high probability conditions, and 80 nontarget trials for low probability conditions. Balanced blocks contained 178 trials and lasted approximately 7.1 minutes each. Cued blocks contained between 177-179 trials and lasted approximately 7.2 minutes each. Total testing time (excluding breaks) was 71.4 minutes.

### 2.4 EEG recording and data processing

EEG was recorded from 128 active electrodes using a Biosemi Active Two system (Biosemi, the Netherlands). Recordings were grounded using common mode sense and driven right leg electrodes (http://www.biosemi.com/faq/cms&drl.htm). Eight channels were added: two electrodes placed 1cm from the outer canthi of each eye, four electrodes placed above and below each eye, and two electrodes placed on the left and right mastoids. EEG was recorded at 1024Hz (DC-coupled with an anti-aliasing filter, −3dB at 204Hz). Electrode offsets were kept within ±50*μ*V.

EEG data were processed using EEGLab V.13.4.4b (Delorme and Makeig, 2004) and ERPLab V.4.0.3.1 (Lopez-Calderon and Luck, 2014) running in MATLAB (The Mathworks). EEG data were downsampled to 512Hz offline. A photosensor was used to measure the timing delay of the video system (10ms) and stimulus event codes were shifted offline to account for this delay. 50Hz line noise was identified using Cleanline (Mullen, 2012) using a separate 1Hz high-pass filtered dataset (EEGLab Basic FIR Filter New, zero-phase, finite impulse response, −6dB cutoff frequency 0.5Hz, transition bandwidth 1Hz). Identified line noise was subtracted from the unfiltered dataset (as recommended by Bigdely-Shamlo et al., 2015). Excessively noisy channels were identified by visual inspection (mean noisy channels by participant = 1.4, median 1, range 0-4) and were excluded from average referencing and independent components analysis (ICA). Data was then referenced to the average of the 128 scalp channels. One channel (FCz) was removed to correct for the rank deficiency caused by average referencing. A separate dataset was processed in the same way, except a 1Hz high-pass filter was applied (filter settings as above) to improve stationarity for the ICA. ICA was performed on the 1Hz high-pass filtered dataset (RunICA extended algorithm, Jung et al., 2000). Independent component information was transferred to the unfiltered dataset. Independent components associated with ocular and muscle activity were identified and removed according to guidelines in Chaumon et al. (2015). Noisy channels and FCz were then interpolated from the cleaned data (spherical spline interpolation). EEG data were low-pass filtered at 30Hz (EEGLab Basic Finite Impulse Response Filter New, zero-phase, −6dB cutoff frequency 33.75Hz, transition band width 7.5Hz). Data were epoched from −100ms to 800ms from test stimulus onset and baseline-corrected using the prestimulus interval. Epochs containing ±100*μ*V deviations from baseline and nontarget trials containing button press responses were rejected.

### 2.5 Statistical Analyses

ERPs were analysed at each electrode and time point using mass univariate repeated measures ANOVAs implemented in the LIMO EEG toolbox v1.4 (Pernet et al., 2011). All time points between −100 and 800ms at all 128 scalp electrodes were included in each analysis (59,008 comparisons). Corrections for multiple comparisons were performed using spatiotemporal cluster corrections based on the cluster mass statistic (Bullmore et al., 1999; Maris & Oostenveld, 2007). 2 x 3 x 2 repeated measures ANOVAs with the factors block (AB/AX), expectation (expected/neutral/surprise) and repetition (repeated/alternating) were performed using the original data and 1000 bootstrap samples. For each bootstrap sample data from both conditions were mean-centred, pooled and then sampled with replacement and randomly allocated to each condition (bootstrap-t method). For each bootstrap sample, all F ratios corresponding to uncorrected p-values of <0.05 were formed into clusters with any neighbouring such F ratios. Channels considered spatial neighbours were defined using the 128-channel Biosemi channel neighbourhood matrix in the LIMO EEG toolbox (Pernet et al., 2011; 2015). Adjacent time points were considered temporal neighbours. The sum of the F ratios in each cluster is the ‘mass’ of that cluster. The most extreme cluster masses in each of the 1000 bootstrap samples were used to estimate the distribution of the null hypothesis. Cluster masses of each cluster identified in the original dataset were compared to the null distribution; the percentile ranking of each cluster relative to the null distribution was used to derive its p-value. The p-value of each cluster was assigned to all members of that cluster. Channel/timepoint combinations not included in any statistically significant cluster were assigned a p-value of 1. These cluster-based multiple comparisons corrections were used because they provide control over the weak family-wise error rate while maintaining high sensitivity to detect broadly-distributed effects (Maris & Oostenveld, 2007; Groppe et al., 2011).

To isolate repetition probability expectation effects signalled by adapter faces in cued blocks, 2 x 2 x 2 repeated measures ANOVAs without neutral conditions were performed with cluster-based corrections as described above.

## 3. Results

### 3.1 Task Performance

Accuracy for detecting and responding to targets collapsed across conditions was high (mean accuracy 97%, range 77-100%). Mean reaction time collapsed across conditions was 470±60ms (range 358-654ms). Accuracy and reaction times were not compared across conditions. This was because of the low number of targets allocated to each condition, and that the task was designed to maintain attention towards nontarget stimuli.

### 3.2 Mass Univariate Analyses of ERPs

### 3.2.1 Main Effects of Repetition

Mass univariate analyses comparing repeated and alternating stimuli revealed repetition effects spanning 99-800ms from test stimulus onset (displayed in Figure 2A). Grand average ERP waveforms evoked by repeated and alternating stimuli at example electrodes are displayed in Figure 2B.

**Figure 2.**
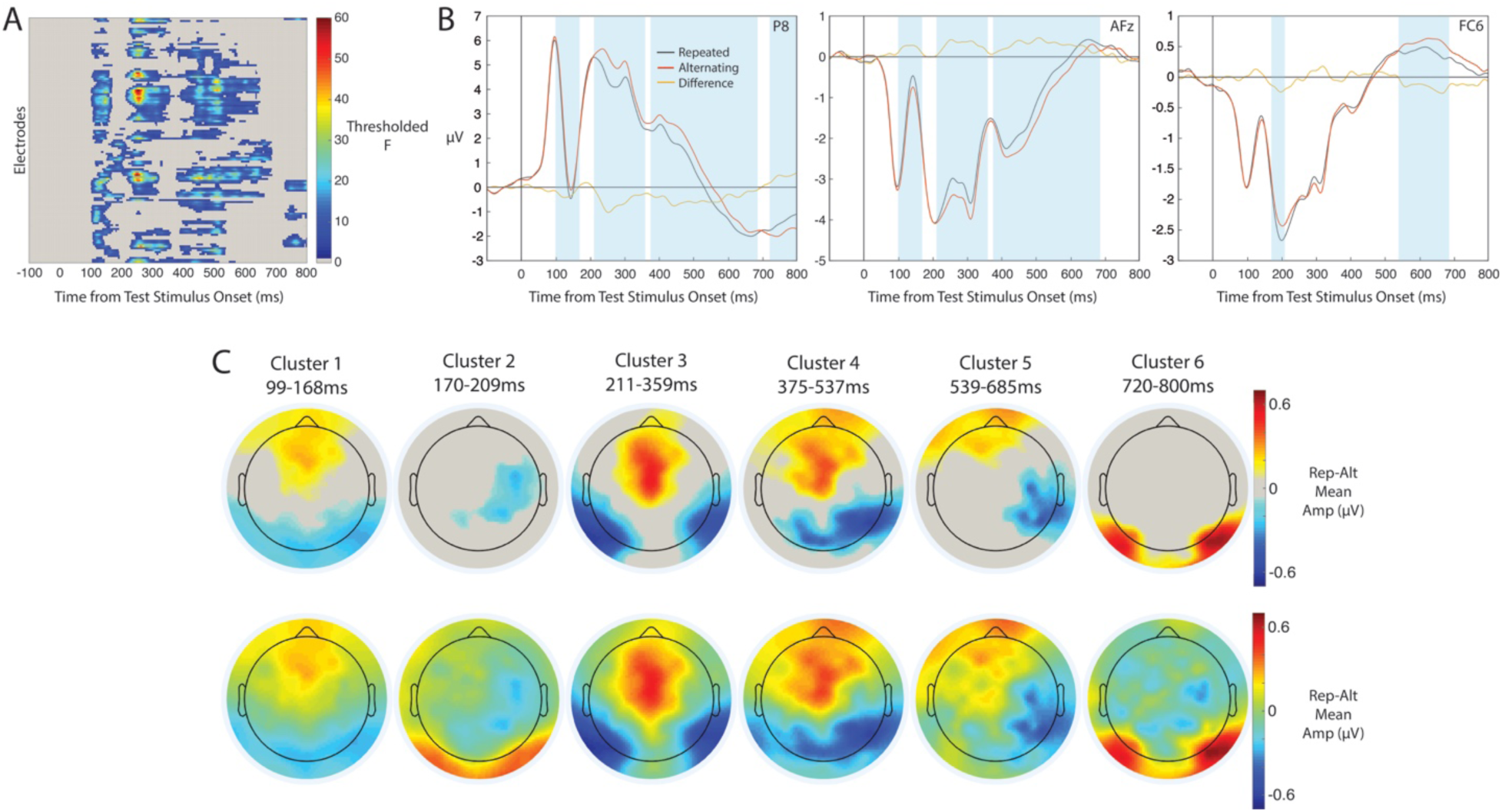
Results of mass univariate repetition effect analyses. A) Timepoint-by-channel matrix of repetition effects. F ratios are displayed thresholded by cluster-level statistical significance. B) Grand-average ERPs evoked by repeated and alternating stimuli. Shaded areas correspond to time windows of repetition effects defined in Figure 2B, for electrodes showing significant effects within each time window. C) Scalp maps of repetition effects by latency from test stimulus onset. Each scalp map shows the mean [repetition - alternation] amplitude difference over the time window, for channel/timepoint combinations included in statistically significant clusters (top row) and all channels (bottom row). (two-column fitting image)

Repetition effects could be broadly divided into 6 time windows (shown in Figure 2C). During the first time window (99-168ms) repeated stimuli evoked more negative-going waveforms at posterior electrodes and more positive waveforms at frontal channels. During the second time window (170-209ms) repeated stimuli evoked more negative-going waveforms at right parietal electrodes. During the third time window (211-359ms) repeated stimuli evoked more negative-going waveforms over bilateral occipitotemporal sites, and more positive-going waveforms over frontocentral channels. During the fourth time window (375-537ms) repeated stimuli evoked more negative-going waveforms at posterior channels, and more positive-going waveforms at frontal and frontocentral channels. During the fifth time window (539-685ms) ERPs were more negative to repeated stimuli at right occipiotemporal channels, and more positive at central and left frontal channels. During the sixth time window (720-800ms) repeated stimuli evoked more positive-going waveforms over bilateral posterior electrodes.

### 3.2.2 Main Effects of Expectation

Comparisons of expected, neutral and surprising stimuli revealed main effects of expectation at time points spanning 113-703ms from test stimulus onset (Figure 3A) which partly overlapped in time with main effects of repetition (Figure 3B). Grand average ERPs to expected, neutral and surprising stimuli at example electrodes are displayed in Figure 3D. Expectation effects could be broadly split into 5 distinct time windows (displayed in Figure 3C). There was an early (113-144ms) over frontal electrodes, followed by an effect between 152-263ms over bilateral occipital, central and frontal electrodes, a third cluster (275-340ms) at central channels, a fourth cluster (342-433ms) at bilateral occipital, central and right frontocentral channels, and a late (435-703ms) effect over right frontocentral and left occipital electrodes.

**Figure 3.**
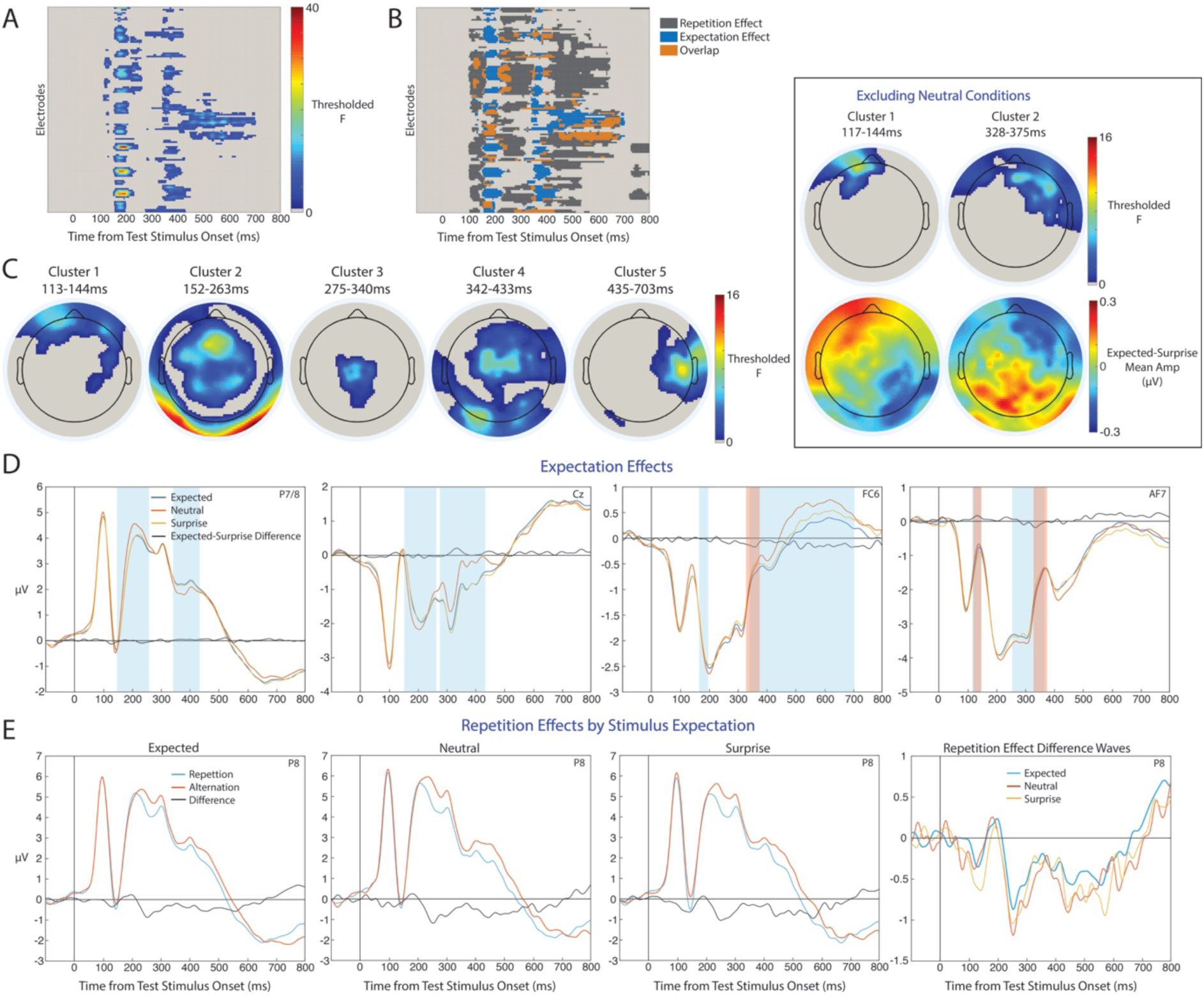
Results of mass univariate expectation effect analyses. A) Timepoint-by-channel matrix of expectation effects. F ratios are thresholded by cluster-level statistical significance. B) Timepoint-by-channel matrix of the spatiotemporal overlap of repetition and expectation effects. C) Scalp maps of expectation effects by latency from test stimulus onset. Average F ratios across each time window are displayed for electrodes within statistically significant clusters. Results of analyses excluding neutral conditions are shown in the boxed area, with [expected – surprising] ERP mean amplitude differences shown under F ratios for each cluster. D) Grand-average ERPs evoked by expected, neutral and surprising test stimuli at selected channels. Shaded areas correspond to time windows of expectation effects defined in Figure 3C, for electrodes showing significant effects within each time window (blue = analyses including neutral conditions, red = analyses excluding neutral conditions). E) Grand-average ERPs showing stimulus repetition effects for expected, neutral and surprising stimuli. (two-column fitting image)

Repeated measures ANOVAs excluding neutral conditions identified 2 clusters of expectation effects (shown in Figure 3C). During the first cluster (117-144ms) expected stimuli evoked more positive-going waveforms at left frontal channels. During the second cluster of effects (328-375ms) ERPs were more negative to expected stimuli over bilateral frontal electrodes.

### 3.2.3 Interactions Involving Expectation and Repetition

ERP repetition effects for expected, neutral and surprising stimuli are displayed at example channel P8 in Figure 3E and were highly similar across the epoch. There were no significant clusters of expectation by repetition interactions, or block by expectation by repetition interactions.

### 3.2.4 Block x Repetition Interactions

There were 3 significant repetition by block interaction clusters between 117-652ms (Figure 4A) which partly overlapped with main effects of repetition (Figure 4B). Topographies of significant interaction clusters are displayed in Figure 4C. Grand-average ERPs evoked by repeated and alternating stimuli in each block type at example channel P8 are shown in Figure 4D. During the first significant interaction cluster (117-179ms) repetition effects were larger in AX compared to AB blocks over bilateral posterior and frontal channels. During the second cluster (246-428ms) the opposite pattern was observed, with larger repetition effects in AB compared to AX blocks. During the third cluster (506-652ms) repetition effects were larger in AX blocks over right occipitotemporal and left frontocentral channels. These interactions were driven by AB/AX block differences for alternating stimulus-evoked ERPs, with no visible block effects on responses to repeated stimuli.

**Figure 4.**
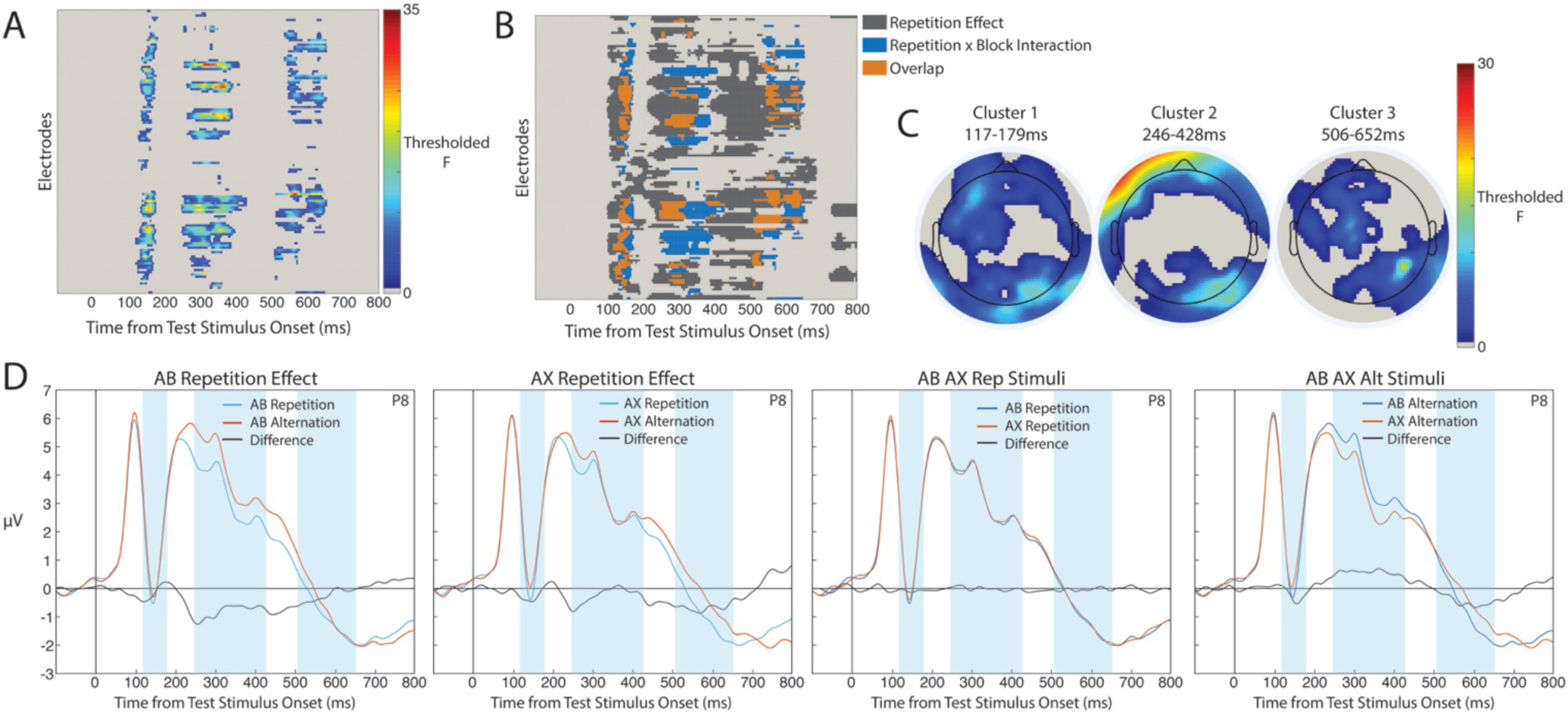
Results of mass univariate block by repetition interaction analyses. A) Timepoint-by-channel matrix of significant repetition by block interaction effects. F ratios are displayed thresholded by cluster-level statistical significance. B) Timepoint-by-channel matrix of the spatiotemporal overlap of statistically significant repetition effects and block by repetition interactions. C) Scalp maps of repetition by block interaction effects by latency from test stimulus onset. Average F ratios across each time window are displayed for electrodes within statistically significant clusters. D) Grand-average ERPs showing repetition effects in AB and AX blocks. Shaded areas correspond to time windows of interaction effects displayed in Figure 4C. (two-column fitting image)

## 4. Discussion

The major finding from this study is that stimulus repetition effects, as measured by ERPs, were not modulated by perceptual expectations. We identified a complex progression of stimulus repetition and expectation effects from 99ms post stimulus onset until the end of the 800ms epoch. Stimulus repetition and expectation effects did partially overlap in time and across electrodes, but did not statistically interact. However, there were differences in the magnitude of repetition effects across blocks of predictable and unpredictable alternating face stimuli. These block differences indicate that repetition effects observed in many previous studies, which presented predictable repeated faces and unpredictable alternating faces, may have conflated effects of repetition and stimulus predictability.

### 4.1 Stimulus Repetition Effects

A range of ERP face image repetition effects could be identified spanning 99-800ms from stimulus onset (displayed in Figure 2) including novel and previously reported effects (reviewed in Schweinberger & Neumann, 2016). An early repetition effect spanned 99-168ms and was similar to previously observed face image repetition effects (e.g. Herzmann et al., 2004; Jemel et al., 2005; but see Caharel et al., 2015). This effect and a later repetition effect spanning 170-209ms from stimulus onset may be due to repetition of low- or mid-level image properties, for example stimulus shape. We also identified the N250r face repetition effect (Schweinberger et al., 2002) spanning 211-359ms from stimulus onset. This effect is larger for face image repetitions (as in our study) compared to when presenting different images of the same face identity (Schweinberger et al., 2002; Caharel et al., 2015; Schweinberger & Neumann, 2016) suggesting that the N250r effect in our data reflects both local and feedforward-inherited effects, characteristic of image repetition effects in high-level visual areas (Vogels, 2016).

Later repetition effects between 375-537ms from stimulus onset differed in direction and topography to previously reported face repetition or semantic/categorical priming effects at similar latencies (e.g. Schweinberger et al., 2002; Wiese & Schweinberger, 2015). These repetition effects may index influences on local recurrent network activity, as found in macaque inferior temporal neurons (Kaliukhovich & Vogels, 2016) and in V1 (Patterson et al., 2013). However, these effects may also index recurrent feedforward and feedback interactions across high-level visual regions (e.g. Ewbank et al., 2011). The late repetition effect from 720-800ms (until the end of the analysed epoch) appears too late to reflect purely local recurrent network activity as reported in Kaliukhovich & Vogels (2016) and may reflect modulation of activity by higher level visual areas, as found in the auditory system (Malmierca et al., 2015).

It is likely that these early and late ERP effects with similar topographies index qualitatively different repetition effects, which may also be associated with distinct changes in stimulus selectivity (e.g. Patterson et al., 2013; Kaliukhovich & Vogels, 2016). This would imply that BOLD repetition effects index a mixture of these early and late repetition effects, which may also be conflated in studies of directed connectivity modulations by stimulus repetition (e.g. de Gardelle et al., 2013; Choi et al., 2017).

### 4.2 Stimulus Expectation Effects

Multiple expectation effects were found spanning 113-703ms from stimulus onset (shown in Figure 3). There were several time periods in which ERPs to neutral stimuli were markedly different to both expected and surprising stimuli, which may partly index effects of block order or time. Block order effects could not be separated in our design, as balanced blocks containing neutral stimuli were always presented before cued blocks containing expected and surprising stimuli. However other experiments controlling for the confound of time have reported smaller BOLD responses to neutral compared with both expected and surprising stimuli (Rahnev et al., 2011; Amado et al., 2016). This suggests that expectation effects, operationalised as stimulus appearance probability, may include different contributions of expectation fulfilment, surprise, and the ability to form expectations for upcoming stimuli (Kovacs & Vogels, 2014; Hsu et al., 2015; Grotheer & Kovacs, 2016).

Analyses were performed excluding neutral blocks in order to isolate effects of stimulus expectations that were cued by adapter stimuli. These analyses revealed an early expectation effect from 117-144ms and a later effect from 328-375ms from stimulus onset. The early effect was statistically significant at frontal channels, however the topography of this effect (shown in Figure 3C) suggests sources in extrastriate visual cortex. This effect topography is consistent with expectation effects based on stimulus transition probabilities in inferior temporal neurons, developed through repeated pairing of stimulus images within trials (Turk-Browne et al., 2009; Meyer & Olson, 2011; Meyer et al., 2014; Ramachandran et al., 2016; Kaposvari et al., 2016). This early effect may be related to similar early (100-200ms) effects of expectations based on repeatedly-paired stimuli in audition (Todorovic & de Lange, 2012).

The later (328-375ms) expectation effect was consistent with frontally-generated effects found when manipulating expectations for abstract stimulus sequences (e.g. expectations of repeated stimulus pairs) that are not associated with any specific stimulus images (e.g. Summerfield et al., 2011). This late effect may correspond to BOLD expectation effects in inferior frontal and middle frontal gyri (Grotheer & Kovacs, 2015; Amado et al., 2016; Choi et al., 2017) and dorsolateral prefrontal cortex (den Ouden et al., 2009; Rahnev et al., 2011). Importantly, these expectation effects are distinct from expectations based on stimulus transition probabilities, and are not found at earlier latencies in extrastriate visual cortex (Kaliukhovich & Vogels, 2011; Meyer et al., 2014; Kaposvari et al., 2016).

The positive dipole effect of the later (328-375ms) expectation effect in our results matches the topography and latency of stimulus expectation effects in the ERP study of Summerfield et al. (2011). It appears that this expectation effect overlapped with the positive dipole of the N250r repetition effect at central electrodes, leading to the appearance of interactive stimulus repetition and expectation effects in their analysis design. This also explains why no expectation effects were identified at bilateral occipitotemporal channels that capture the negative dipole of the N250r repetition effect (Schweinberger et al., 2002; Neumann & Schweinberger, 2016) which was reported in Summerfield et al. to be modulated by expectation.

### 4.3 Modulations of Repetition Effects by Expectation

ERP repetition and expectation effects did not statistically interact throughout the epoch, in agreement with previous findings that stimulus repetition effects are distinct from effects of perceptual expectations (Grotheer & Kovacs, 2015; Kaliukhovich & Vogels, 2011, 2014). Together, these findings support models that specify separate repetition and perceptual expectation effects (e.g. Grotheer & Kovacs, 2016; Grimm et al., 2016; Henson, 2016; Vogels, 2016) and provide evidence against models proposing a modulatory effect of perceptual expectation on the magnitude of repetition suppression (Summerfield et al., 2008, 2011; Auksztukewicz & Friston, 2016). Additionally, studies reporting changes in directed connectivity with stimulus repetition consistent with predictive coding models have used designs that confound stimulus repetition and expectation (e.g. Garrido et al., 2009; Ewbank et al., 2011; de Gardelle et al., 2013; reviewed in Auksztulewicz & Friston, 2016). Importantly, these connectivity changes could also plausibly occur from additive repetition and expectation effects. Modelling results from experiments the orthogonally manipulate repetition and expectation are needed to verify whether repetition effects are associated with changes in top-down connectivity.

It is important to note that we cannot directly support the null hypothesis of no expectation by repetition interaction using our frequentist mass univariate testing approach. Future studies could focus on individual repetition effect time windows, and adopt Bayesian hypothesis testing to quantify evidence for null and alternative hypotheses (Rouder et al., 2009). This could be done using priors informed by our study and previous face repetition experiments (for a review of face repetition experiments see Neumann & Schweinberger, 2016). However, other sources of information support the claim that repetition effects are not a product of perceptual expectations. The main effects of repetition and expectation in our results had differing topographies and only partially overlapped in time (Figure 3B), indicating distinct processes. In addition, any differences in point estimates of repetition effects by expectation appeared to be much smaller than the repetition effects themselves (shown in Figure 3E), contrary to the claim that repetition effects are not found for surprising stimuli (e.g. Summerfield et al., 2008). As our results also explain the reported expectation by repetition interaction in Summerfield et al. (2011) as a mixture of additive effects, it appears that there is currently no electrophysiological evidence of modulations of repetition suppression by expectation, at least for studies using visual stimuli.

Findings of noninteracting repetition and perceptual expectation effects in our study and those of other experiments using electrophysiological recordings (Kaliukhovich & Vogels, 2011; 2014) contrast with reports of larger BOLD repetition effects for surprising stimuli (e.g. Amado et al., 2016; de Gardelle et al., 2013; Larsson & Smith, 2012; reviewed in Kovacs & Vogels, 2014). One possibility is that these BOLD increases to surprising alternations result from increased gamma-band activity (e.g. Todorovic et al., 2011), hypothesised to signal feedforward prediction errors in supragranular cortical layers (Auksztulewicz & Friston, 2016). Gamma-band activity can be closely coupled with the BOLD response (Niessing et al., 2005) but would not be found in the ERPs analysed in our study. Future studies using MEG, electrocorticography or intracranial electrodes will be able to determine whether surprise responses, as reflected in gamma band activity, are suppressed by stimulus repetition.

### 4.4 Differences in Repetition Effects by Block Type

Magnitudes of repetition effects differed across AB and AX block types, with stimuli in each block type showing larger repetition effects at distinct latencies from stimulus onset. Block differences were found specifically for alternating stimuli, indicating that block effects did not modulate the underlying repetition effect mechanisms, but rather affected the observed repetition/alternation signal difference (see Feuerriegel, 2016 for further discussion of this issue). Our findings indicate that many previous experiments, which have used predictable repeated stimuli and unpredictable alternating stimuli, may have indexed stimulus predictability effects as part of the observed repetition effect.

One limitation of this study is that we could not dissociate novelty and predictability effects in our experimental design. Larger repetition effects in AX blocks likely index effects of both increased novelty for AX alternating stimuli (e.g. Xiang & Brown, 1998; Mur et al., 2010) and the inability to form image-specific expectations for alternating stimulus images in AX blocks (e.g. Hsu et al., 2014, 2015, 2016). Future research controlling for stimulus novelty may be able to identify and isolate stimulus feature predictability effects in repetition designs, which have received little attention in the repetition suppression literature (but see Pajani et al., 2017; Feuerriegel, 2016).

It is unclear why AB blocks showed larger repetition effects between 246-428ms (during the N250r repetition effect). One possibility is that population responses to the often-presented alternating stimuli in AB blocks became more distinct from those to other faces in inferior temporal areas through repeated stimulus presentation (familiarisation; e.g. Freedman et al., 2006; Meyer et al., 2014b). More distinct population response patterns between faces would lead to less cross-stimulus adaptation of alternating stimuli by visually-similar adapters (Verhoef et al., 2008; De Baene & Vogels, 2010) resulting in larger differences between repeated and alternating stimulus-evoked ERPs.

### 4.5 Conclusions

This research has systematically, and using a data driven approach, identified spatiotemporal sequences of separable repetition and expectation effects on ERPs in the one experimental paradigm. Our results support two-stage models of repetition suppression that pose a distinction between repetition effects driven by prior stimulus exposure and top-down perceptual expectations (Grotheer & Kovacs, 2016; Grimm et al., 2016; Henson, 2016; Vogels, 2016).

## Acknowledgements

We thank Dilushi Chandrakumar for her assistance with preparation of stimuli. Daniel Feuerriegel was supported by an Australian Government Research Training Program Scholarship. Funding sources had no role in study design, data collection, analysis or interpretation of results.

## References

Amado, C., Hermann, P., Kovacs, P., Grotheer, M., Vidnyanszky, Z., & Kovacs, G. (2016). The contribution of surprise to the prediction based modulation of fMRI responses. Neuropsychologia, 84, 105–112.

Auksztulewicz, R., & Friston, K. (2016). Repetition suppression and its contextual determinants in predictive coding. Cortex, 80, 125–140.

Bigdely-Shamlo, N., Mullen, T., Kothe, C., Su, K. M., & Robbins, K. A. (2015). The PREP pipeline: standardized preprocessing for large-scale EEG analysis. Frontiers in Neuroinformatics, 9, 16.

Brainard, D. H. (1997). The Psychophysics Toolbox. Spatial Vision, 10(4), 433–436.

Bullmore, E. T., Suckling, J., Overmeyer, S., Rabe-Hesketh, S., Taylor, E., & Brammer, M. J. (1999). Global, voxel, and cluster tests, by theory and permutation, for a difference between two groups of structural MR images of the brain. IEEE Transactions on Medical Imaging, 18(1), 32–42.

Caharel, S., Collet, K., & Rossion, B. (2015). The early visual encoding of a face (N170) is viewpoint-dependent: a parametric ERP-adaptation study. Biological Psychology, 106, 18–27.

Chaumon, M., Bishop, D. V., & Busch, N. A. (2015). A practical guide to the selection of independent components of the electroencephalogram for artifact correction. Journal of Neuroscience Methods, 250, 47–63.

Choi, U.-S., Sung, Y.-W., & Ogawa, S. (2017). Steady-state and dynamic network modes for perceptual expectation. Scientific Reports, 7.

De Baene, W., & Vogels, R. (2010). Effects of adaptation on the stimulus selectivity of macaque inferior temporal spiking activity and local field potentials. Cerebral Cortex, 20(9), 2145–2165.

de Gardelle, V., Waszczuk, M., Egner, T., & Summerfield, C. (2013). Concurrent repetition enhancement and suppression responses in extrastriate visual cortex. Cerebral Cortex, 23(9), 2235–2244.

Delorme, A., & Makeig, S. (2004). EEGLAB: an open source toolbox for analysis of single-trial EEG dynamics including independent component analysis. Journal of Neuroscience Methods, 134(1), 9–21.

Den Ouden, H. E., Friston, K. J., Daw, N. D., McIntosh, A. R., & Stephan, K. E. (2009). A dual role for prediction error in associative learning. Cerebral Cortex, 19(5), 1175–1185.

Desimone, R. (1996). Neural mechanisms for visual memory and their role in attention. Proceedings of the National Academy of Sciences of the United States of America, 93(24), 13494–13499.

Dhruv, N. T., & Carandini, M. (2014). Cascaded effects of spatial adaptation in the early visual system. Neuron, 81(3), 529–535.

Ebner, N. C. (2008). Age of face matters: Age-group differences in ratings of young and old faces. Behavior Research Methods, 40(1), 130–136.

Egner, T., Monti, J. M., & Summerfield, C. (2010). Expectation and surprise determine neural population responses in the ventral visual stream. Journal of Neuroscience, 30(49), 16601–16608.

Ewbank, M. P., Lawson, R. P., Henson, R. N., Rowe, J. B., Passamonti, L., & Calder, A. J. (2011). Changes in “top-down” connectivity underlie repetition suppression in the ventral visual pathway. Journal of Neuroscience, 31(15), 5635–5642.

Feuerriegel, D. (2016). Selecting appropriate designs and comparison conditions in repetition paradigms. Cortex, 80, 196–205.

Freedman, D. J., Riesenhuber, M., Poggio, T., & Miller, E. K. (2006). Experience-dependent sharpening of visual shape selectivity in inferior temporal cortex. Cerebral Cortex, 16(11), 1631–1644.

Friston, K. (2005). A theory of cortical responses. Philosophical Transactions of the Royal Society of London. Series B: Biological Sciences, 360(1456), 815–836.

Friston, K., & Kiebel, S. (2009). Predictive coding under the free-energy principle. Philosophical Transactions of the Royal Society of London. Series B: Biological Sciences, 364(1521), 1211–1221.

Garrido, M. I., Kilner, J. M., Kiebel, S. J., Stephan, K. E., Baldeweg, T., & Friston, K. J. (2009). Repetition suppression and plasticity in the human brain. Neuroimage, 48(1), 269–279.

Grill-Spector, K., Henson, R., & Martin, A. (2006). Repetition and the brain: Neural models of stimulus-specific effects. Trends in Cognitive Science, 10(1), 14–23.

Grimm, S., Escera, C., & Nelken, I. (2016). Early indices of deviance detection in humans and animal models. Biological Psychology, 116, 23–27.

Groppe, D. M., Urbach, T. P., & Kutas, M. (2011). Mass univariate analysis of eventrelated brain potentials/fields I: a critical tutorial review. Psychophysiology, 48(12), 1711–1725.

Grotheer, M., & Kovacs, G. (2014). Repetition probability effects depend on prior experiences. Journal of Neuroscience, 34(19), 6640–6646.

Grotheer, M., & Kovacs, G. (2015). The relationship between stimulus repetitions and fulfilled expectations. Neuropsychologia, 67, 175–182.

Grotheer, M., & Kovacs, G. (2016). Can predictive coding explain repetition suppression? Cortex, 80, 113–124.

Herzmann, G., Schweinberger, S. R., Sommer, W., & Jentzsch, I. (2004). What's special about personally familiar faces? A multimodal approach. Psychophysiology, 41(5), 688–701.

Hsu, Y. F., Hamalainen, J. A., & Waszak, F. (2013). Temporal expectation and spectral expectation operate in distinct fashion on neuronal populations. Neuropsychologia, 51(13), 2548–2555. doi:10.1016/j.neuropsychologia.2013.09.01.

Hsu, Y. F., Hamalainen, J. A., & Waszak, F. (2014). Both attention and prediction are necessary for adaptive neuronal tuning in sensory processing. Frontiers in Human Neuroscience, 8, 152.

Hsu, Y. F., Le Bars, S., Hamalainen, J. A., & Waszak, F. (2015). Distinctive Representation of Mispredicted and Unpredicted Prediction Errors in Human Electroencephalography. Journal of Neuroscience, 35(43), 14653–14660.

Jemel, B., Pisani, M., Rousselle, L., Crommelinck, M., & Bruyer, R. (2005). Exploring the functional architecture of person recognition system with event-related potentials in a within-and cross-domain self-priming of faces. Neuropsychologia, 43(14), 2024–2040. doi:10.1016/j.neuropsychologia.2005.03.01.

Jung, T. P., Makeig, S., Westerfield, M., Townsend, J., Courchesne, E., & Sejnowski, T. J. (2000). Removal of eye activity artifacts from visual event-related potentials in normal and clinical subjects. Clinical Neurophysiology, 111(10), 1745–1758.

Kaliukhovich, D. A., & Vogels, R. (2011). Stimulus repetition probability does not affect repetition suppression in macaque inferior temporal cortex. Cerebral Cortex, 21(7), 1547–1558.

Kaliukhovich, D. A., & Vogels, R. (2014). Neurons in macaque inferior temporal cortex show no surprise response to deviants in visual oddball sequences. Journal of Neuroscience, 34(38), 12801–12815.

Kaliukhovich, D. A., & Vogels, R. (2016). Divisive normalization predicts adaptation-induced response changes in Macaque inferior temporal cortex. Journal of Neuroscience, 36(22), 6116–6128.

Kanwisher, N., McDermott, J., & Chun, M. M. (1997). The fusiform face area: A module in human extrastriate cortex specialized for face perception. Journal of Neuroscience, 17(11), 4302–4311.

Kaposvari, P., Kumar, S., & Vogels, R. (2016). Statistical Learning Signals in Macaque Inferior Temporal Cortex. Cerebral Cortex.

Kleiner, M., Brainard, D., Pelli, D., Ingling, A., Murray, R., & Broussard, C. (2007). What's new in Psychtoolbox-3. Perception, 36(14), 1.

Kohn, A. (2007). Visual adaptation: physiology, mechanisms, and functional benefits. Journal of Neurophysiology, 97(5), 3155–3164.

Kovacs, G., Iffland, L., Vidnyanszky, Z., & Greenlee, M. W. (2012). Stimulus repetition probability effects on repetition suppression are position invariant for faces. Neuroimage, 60(4), 2128–2135.

Kovacs, G., Kaiser, D., Kaliukhovich, D. A., Vidnyanszky, Z., & Vogels, R. (2013). Repetition probability does not affect fMRI repetition suppression for objects. Journal of Neuroscience, 33(23), 9805–9812.

Kovacs, G., & Vogels, R. (2014). When does repetition suppression depend on repetition probability? Frontiers in Human Neuroscience, 8, 685.

Larsson, J., & Smith, A. T. (2012). fMRI repetition suppression: neuronal adaptation or stimulus expectation? Cerebral Cortex, 22(3), 567–576.

Larsson, J., Solomon, S. G., & Kohn, A. (2016). fMRI adaptation revisited. Cortex, 80, 154–160.

Lopez-Calderon, J., & Luck, S. J. (2014). ERPLAB: an open-source toolbox for the analysis of event-related potentials. Frontiers in Human Neuroscience, 8, 213.

Malmierca, M. S., Anderson, L. A., & Antunes, F. M. (2015). The cortical modulation of stimulus-specific adaptation in the auditory midbrain and thalamus: A potential neuronal correlate for predictive coding. Frontiers in Systems Neuroscience, 9, 19.

Maris, E., & Oostenveld, R. (2007). Nonparametric statistical testing of EEG-and MEG-data. Journal of Neuroscience Methods, 164(1), 177–190.

Meyer, T., & Olson, C. R. (2011). Statistical learning of visual transitions in monkey inferotemporal cortex. Proceedings of the National Academy of Sciences of the United States of America, 108(48), 19401–19406.

Meyer, T., Ramachandran, S., & Olson, C. R. (2014). Statistical learning of serial visual transitions by neurons in monkey inferotemporal cortex. Journal of Neuroscience, 34(28), 9332–9337.

Meyer, T., Walker, C., Cho, R. Y., & Olson, C. R. (2014). Image familiarization sharpens response dynamics of neurons in inferotemporal cortex. Nature Neuroscience, 17(10), 1388–1394.

Minear, M., & Park, D. C. (2004). A lifespan database of adult facial stimuli. Behavior Research Methods, Instruments, & Computers, 36(4), 630–633.

Movshon, J. A., & Lennie, P. (1979). Pattern-selective adaptation in visual cortical neurones. Nature, 278(5707), 850–852.

Mullen, T. (2012). CleanLine EEGLAB plugin. San Diego, CA: Neuroimaging Informatics Tools and Resources Clearinghouse (NITRC).

Mur, M., Ruff, D. A., Bodurka, J., Bandettini, P. A., & Kriegeskorte, N. (2010). Face-identity change activation outside the face system: “Release from adaptation” may not always indicate neuronal selectivity. Cerebral Cortex, 20(9), 2027–2042.

Nicholls, M. E., Thomas, N. A., Loetscher, T., & Grimshaw, G. M. (2013). The Flinders Handedness survey (FLANDERS): a brief measure of skilled hand preference. Cortex, 49(10), 2914–2926.

Niessing, J., Ebisch, B., Schmidt, K. E., Niessing, M., Singer, W., & Galuske, R. A. (2005). Hemodynamic signals correlate tightly with synchronized gamma oscillations. Science, 309(5736), 948–951.

Pajani, A., Kouider, S., Roux, P., & de Gardelle, V. (2017). Unsuppressible repetition suppression and exemplar-specific expectation suppression in the fusiform face area. Scientific Reports, 7, 160.

Patterson, C. A., Wissig, S. C., & Kohn, A. (2013). Distinct effects of brief and prolonged adaptation on orientation tuning in primary visual cortex. Journal of Neuroscience, 33(2), 532–543.

Pernet, C. R., Chauveau, N., Gaspar, C., & Rousselet, G. A. (2011). LIMO EEG: A toolbox for hierarchical LInear MOdeling of ElectroEncephaloGraphic data. Computational Intelligence and Neuroscience, 2011, 831409.

Pernet, C. R., Latinus, M., Nichols, T. E., & Rousselet, G. A. (2015). Cluster-based computational methods for mass univariate analyses of event-related brain potentials/fields: A simulation study. Journal of Neuroscience Methods, 250, 85–93.

Rahnev, D., Lau, H., & de Lange, F. P. (2011). Prior expectation modulates the interaction between sensory and prefrontal regions in the human brain. Journal of Neuroscience, 31(29), 10741–10748.

Rao, R. P., & Ballard, D. H. (1999). Predictive coding in the visual cortex: A functional interpretation of some extra-classical receptive-field effects. Nature Neuroscience, 2(1), 79–87.

Rouder, J. N., Speckman, P. L., Sun, D., Morey, R. D., & Iverson, G. (2009). Bayesian t tests for accepting and rejecting the null hypothesis. Psychonomic Bulletin and Review, 16(2), 225–237.

Sawamura, H., Orban, G. A., & Vogels, R. (2006). Selectivity of neuronal adaptation does not match response selectivity: A single-cell study of the FMRI adaptation paradigm. Neuron, 49(2), 307–318.

Schweinberger, S. R., & Neumann, M. F. (2016). Repetition effects in human ERPs to faces. Cortex, 80, 141–153.

Schweinberger, S. R., Pickering, E. C., Jentzsch, I., Burton, A. M., & Kaufmann, J. M. (2002). Event-related brain potential evidence for a response of inferior temporal cortex to familiar face repetitions. Brain Research: Cognitive Brain Research, 14(3), 398–409.

Segaert, K., Weber, K., de Lange, F. P., Petersson, K. M., & Hagoort, P. (2013). The suppression of repetition enhancement: A review of fMRI studies. Neuropsychologia, 51(1), 59–66.

Solomon, S. G., & Kohn, A. (2014). Moving sensory adaptation beyond suppressive effects in single neurons. Current Biology, 24(20), R1012–1022.

Summerfield, C., Trittschuh, E. H., Monti, J. M., Mesulam, M. M., & Egner, T. (2008). Neural repetition suppression reflects fulfilled perceptual expectations. Nature Neuroscience, 11(9), 1004–1006.

Summerfield, C., Wyart, V., Johnen, V. M., & de Gardelle, V. (2011). Human scalp electroencephalography reveals that repetition suppression varies with expectation. Frontiers in Human Neuroscience, 5, 67.

Todorovic, A., & de Lange, F. P. (2012). Repetition suppression and expectation suppression are dissociable in time in early auditory evoked fields. Journal of Neuroscience, 32(39), 13389–13395.

Todorovic, A., van Ede, F., Maris, E., & de Lange, F. P. (2011). Prior expectation mediates neural adaptation to repeated sounds in the auditory cortex: An MEG study. Journal of Neuroscience, 31(25), 9118–9123.

Tottenham, N., Tanaka, J. W., Leon, A. C., McCarry, T., Nurse, M., Hare, T. A., … Nelson, C. (2009). The NimStim set of facial expressions: Judgments from untrained research participants. Psychiatry Research, 168(3), 242–249.

Turk-Browne, N. B., Scholl, B. J., Chun, M. M., & Johnson, M. K. (2009). Neural evidence of statistical learning: Efficient detection of visual regularities without awareness. Journal of Cognitive Neuroscience, 21(10), 1934–1945.

Verhoef, B. E., Kayaert, G., Franko, E., Vangeneugden, J., & Vogels, R. (2008). Stimulus similarity-contingent neural adaptation can be time and cortical area dependent. Journal of Neuroscience, 28(42), 10631–10640.

Vogels, R. (2016). Sources of adaptation of inferior temporal cortical responses. Cortex, 80, 185–195.

Whitmire, C. J., & Stanley, G. B. (2016). Rapid sensory adaptation redux: A circuit perspective. Neuron, 92(2), 298–315.

Wiese, H., & Schweinberger, S. R. (2015). Getting connected: Both associative and semantic links structure semantic memory for newly learned persons. Quarterly Journal of Experimental Psychology (2006), 68(11), 2131–2148.

Willenbockel, V., Sadr, J., Fiset, D., Horne, G. O., Gosselin, F., & Tanaka, J. W. (2010). Controlling low-level image properties: The SHINE toolbox. Behavior Research Methods, 42(3), 671–684.

Wissig, S. C., & Kohn, A. (2012). The influence of surround suppression on adaptation effects in primary visual cortex. Journal of Neurophysiology, 107(12), 3370–3384.

Xiang, J. Z., & Brown, M. W. (1998). Differential neuronal encoding of novelty, familiarity and recency in regions of the anterior temporal lobe. Neuropharmacology, 37(4-5), 657–676.

